# Evaluating Approaches for Inference Testing of Whole-Brain Densely Sampled Single-Subject Task fMRI Data

**DOI:** 10.64898/2026.06.29.735344

**Authors:** Michelle C. Medina, Neha A. Reddy, Molly G. Bright, Kevin R. Sitek

**Author notes:** Corresponding Author: Michelle C. Medina, 645 N. Michigan Ave., Suite 1100, Chicago, IL 60611 USA.

## Abstract

Task-based precision mapping has become a promising technique in functional MRI (fMRI) to robustly characterize and map an individual’s unique activity patterns. These experiments consist of acquiring extensive imaging data in one participant, ultimately improving the sensitivity and specificity of individual-specific functional localization. Despite its advantages, studies have primarily focused on understanding individual-specific cortical activation, preventing a holistic view of a systems-level functional response, and to date, best approaches for the statistical analysis of controlled task-based, densely sampled, whole-brain data have not yet been fully established. Therefore, in this study, we collected whole-brain (i.e. covering cortex, cerebellum, and brainstem) multi-echo densely sampled data of the auditory system, a system with major subcortical components, and evaluated activation sensitivity as well as activation stability across data subsets of commonly-used whole-brain and region-specific inference testing approaches. The whole-brain approaches involved standard voxel-level and cluster-level inference schemes with varying statistical thresholds and a non-parametric permutation inference approach. The region-specific approaches involved an exploratory top % t-statistics methods and non-parametric permutation inference approaches. We found that a whole-brain voxel-level approach with a false discovery rate (FDR) correction (p<0.05) presented highest sensitivity across regions and subjects as well as most consistent detection of expected auditory regions, even with lower scan duration. In addition, we found that a region-specific top % t-statistic approach may be a useful exploratory functional localization tool and a complementary method to standard inference testing approaches.

## 1. Introduction

Precision mapping (Gordon et al., 2017) has become a promising technique in functional MRI (fMRI) to robustly characterize and map an individual’s unique activity patterns. These experiments consist of acquiring extensive imaging data in one participant, which has been shown to improve the sensitivity and specificity of individual-specific functional localization (Gordon et al., 2017; Hemmerling et al., 2025; Medina et al., 2026). Previous studies have employed precision mapping strategies primarily in resting state fMRI experiments (Braga & Buckner, 2017; Gordon et al., 2017; Gratton et al., 2018; Lynch et al., 2020), yielding high fidelity functional connectivity maps and showcasing differences in network organization across densely sampled individuals. Additionally, these studies have closely examined the differences between participant-specific networks and group-averaged networks (Braga & Buckner, 2017; Gordon et al., 2017; Gratton et al., 2018), revealing the existence of individual-specific variations in the spatial locations of functional networks and highlighting the promising advantages of dense sampling approaches in the understanding of functional signatures within individuals, which may be critically beneficial for clinical populations (D’Esposito, 2019; Satterthwaite et al., 2018).

Recently, precision mapping has also been used with task fMRI to functionally map task-based activity, providing insight into individual-specific brain function (Ceja et al., 2024; Gordon et al., 2017; Lynch et al., 2020, 2021; Medina et al., 2026). Similar to the findings in resting state studies, results show added benefits in using densely sampled data with more robust localization of individual-specific task-based functional activation compared to group-averaged responses, which may be sparse and challenging to register to a standard brain (Gordon et al., 2017; Medina et al., 2026). Despite its promising benefits, task-based precision mapping is comparatively underexplored, and its literature presents two major limitations. First, studies have primarily focused on understanding individual-specific cortical activation, which does not provide a holistic view of a systems-level functional response. Recent work in resting state precision functional mapping has begun exploring the functional responses of subcortical and cerebellar regions (Greene et al., 2020; Lynch et al., 2020; Marek et al., 2018; Marek & Greene, 2021), and these investigations are likewise necessary in densely-sampled task fMRI studies to obtain a more comprehensive understanding of entire neural systems. Although subcortical regions present various challenges such as low tSNR (Maugeri et al., 2018), poor contrast (De Martino et al., 2015; Saranathan et al., 2021; Williams et al., 2024), small nuclei (Glendenning & Masterton, 1998; Sitek et al., 2019), and vascular and physiological confounds (Beissner, 2015), recent advancements have made it possible to successfully identify cortical, subcortical, and brainstem task activation in different sensory neural systems by using whole-brain acquisitions (Brooks et al., 2017; Oliva et al., 2022) and multi-echo approaches (Medina et al., 2026; Reddy et al., 2024). Thus, to address this first limitation and contribute an open-source task-based precision mapping dataset to the community, we collected whole-brain (i.e. covering cortex, cerebellum, and brainstem) multi-echo densely sampled data of the auditory system, a system with major subcortical components.

Second, to date, best approaches for the analysis of controlled task-based, densely sampled, whole-brain data have not yet been fully established. Current work exhibits variability in its methodology for activation localization and statistical inference. For instance, although previous precision mapping task fMRI studies have identified functional responses in expected task regions with whole-brain voxel-wise (Gordon et al., 2017) and cluster-corrected (Ceja et al., 2024) analyses, these findings have either not detailed multiple comparison corrections or inflated false positives have been reported in their results, respectively. Therefore, it remains uncertain which methods may provide more robust and reliable results when using precision mapping task-based datasets.

There exists a great body of work that has focused on evaluating conventional inference methods in traditional single-subject or group-level fMRI data. For example, previous studies have applied different voxel-level thresholding techniques in simulated (Logan & Rowe, 2004; Marchini & Presanis, 2004) and publicly available (Soltysik, 2020) single-subject fMRI data, showing that a false discovery rate (FDR) may provide less stringent thresholding than family-wise error (FWE) rate corrections (Lindquist & Mejia, 2015) and thus may provide a higher power in the detection of activated voxels (Benjamini & Hochberg, 1995; Logan & Rowe, 2004). Cluster-level thresholding with FWE rate correction has also become a popular approach, especially at the group-level (Hausfeld et al., 2024; Medina et al., 2026; Reddy et al., 2024; Woo et al., 2014), due to its increased sensitivity and consideration of the spatial correlation between fMRI voxels (Friston et al., 1994; Smith & Nichols, 2009; Woo et al., 2014), but comparisons with other methods have showcased its limitations with spatial specificity (Lindquist & Mejia, 2015; Soltysik, 2020; Woo et al., 2014). Recently, non-parametric permutation inference methods (Nichols & Holmes, 2001) have shown promising results in traditional group-level task fMRI data, particularly when paired with region-specific analyses, which better elucidate activation in specific regions that are known *a priori* to be involved with the examined functional task (Medina et al., 2026; Oliva et al., 2022; Reddy et al., 2024). Additionally, region-specific qualitative thresholds, such as identifying voxels with the top percentile of t-statistics, have been successfully employed to delineate functionally active regions in precision mapping data (Medina et al., 2026; Salvo, Anderson, et al., 2025; Salvo, Lakshman, et al., 2025). However, given that these methods have not been evaluated in whole-brain densely sampled single-subject data, it is difficult to determine best practices and understand their limitations, especially when exploring activation across subcortical regions where sensitivity and detection of significant functional activation may be challenging with conventional inference approaches. Thus, to address this second limitation, we performed a comparative assessment of standard whole-brain (voxel-level, cluster-level, and non-parametric permutation) and region-specific (top % of t-statistics and non-parametric permutation) inference methods by evaluating (1) their ability to identify significant activation in expected auditory regions at varying standard thresholds and (2) the reliability of spatial localization and spatial overlap of the resulting activated clusters within the same whole-brain precision mapping dataset.

Ultimately, task-based precision mapping fMRI is a promising tool to better understand individual-specific brain function for basic neuroscience and clinical utility. However, to achieve reliable precision mapping, we need to be able to robustly capture whole-brain information and understand best practices for activity localization and statistical inference. Therefore, in this study, we collect whole-brain densely sampled single-subject data and investigate the capacity of commonly used inference approaches to reliably identify regions of activation and characterize significance across the brain in a well-known sensory system.

## 2. Methods

### 2.1 Data Collection

Five healthy individuals (25±1y, 3F) with no known history of neurological, vascular, or auditory disorders were enrolled in the study after providing written, informed consent. Data from three of the individuals (i.e. sub-01, sub-02, and sub-03) were collected as part of a previous study (Medina et al., 2026). The study protocol was approved by the Institutional Review Board of Northwestern University. Participants were scanned using a Siemens 3T Prisma and 32-channel head coil. A multi-echo T1-weighted MPRAGE was collected and used for registration (Tisdall et al., 2016): TR = 2.17 s, TEs = 1.69/3.55/5.41 ms, TI = 1.16 s, FA = 7°, voxel size = 1 x 1 x 1 mm^3^, FOV = 256 x 256 m^2^. The three echo MPRAGE images were then combined using root-mean-square. Four functional scans (10 minutes of functional data per scan) were collected using a multi-band, multi-echo gradient-echo echo-planar imaging sequence with whole-brain coverage provided by the Center for Magnetic Resonance Research (CMRR, Minnesota) (Moeller et al., 2010; Setsompop et al., 2012): TR=2.2s, TEs=13.40/39.5/65.6ms, MB factor=2, GRAPPA phase-encoding acceleration factor=2, voxel size=1.731×1.731×4mm^3^, 44 slices, FOV=180mm, and 250 volumes (Moeller et al., 2010; Reddy et al., 2024; Setsompop et al., 2012). Axial slices were aligned perpendicular to the base of the fourth ventricle to align with brainstem anatomy while achieving maximum whole-brain coverage. A reverse phase encode functional image was collected and used for distortion correction (Beissner, 2015). During all scanning, end-tidal CO_2_ was measured using a nasal cannula and gas analyzer system at a sampling rate of 1000 Hz (PowerLab, ADInstruments).

### 2.2 Auditory Setup and Stimulus

Avotec’s Silent Scan system and Sensimetric S14 earbuds with disposable foam canal ear tips (Comply) were used to transmit the audio with an average noise reduction rating above 29 dB, as per the manufacturer’s advertised specifications. MULTIPAD EAR inflatable pads (Pearl Technology) were placed between the head coil and the patient for additional comfort and immobilization. Participants were instructed to attentively listen to the music, remain as still as possible, and stay awake while looking at a fixation cross. During each functional scan, participants listened to the auditory stimuli in a block design: 10-seconds of an RMS-normalized, instrumental-only pop song was followed by 15-seconds rest for a total of 21 trials. Three different pop songs were used to maintain attention, and song choice was randomized across participants and scans.

### 2.3 Data Analysis

#### 2.3.1 MRI Pre-processing

Data were analyzed with similar methods to those used in Medina et al (Medina et al., 2026). All pre-processing steps were performed using FSL (Jenkinson et al., 2012) (version 6.0.3) and AFNI (Cox, 1996) (version 24.1.11). T1-weighted images for all participants were processed, including bias field correction and brain extraction (*fsl_anat*). The first 10 volumes of each functional scan were removed for the signal to reach steady-state equilibrium. Motion realignment parameters were calculated using the first echo data and the Single Band reference image (*3dvolreg*) and applied to all echoes (*3dAllineate*). Brain extraction (*bet*) and distortion correction (*topup, applytopup*) were performed on all images.

#### 2.3.2 Multi-echo with independent component analysis (ME-ICA)

Echo timeseries were optimally combined (DuPre et al., 2019) and ICA decomposition was performed using tedana (DuPre et al., 2021) (version 24.0.2). Rejected (i.e., noise-related) ICA components were automatically classified using tedana’s external regressors decision tree (*decision_tree_demo_external_regressors_motion_task_models*) (Community et al., 2024), which included a cerebrospinal fluid regressor, six motion regressors and their derivatives as nuisance regressors. The ICA components were then orthogonalized using the conservative approach in Moia et al (Moia et al., 2021) and Reddy et al (Reddy et al., 2025). The optimally combined data were smoothed using a 3 mm FWHM Gaussian blur (*3dMerge*) and converted to signal percent change.

#### 2.3.4 General Linear Model

Increased amounts of individual-subject data have resulted in more reliable estimates of individual-subject activation and an increase in individual functional specificity (Gordon et al., 2017; Medina et al., 2026). Thus, to generate the densely sampled data, all four functional scans were concatenated prior to subject-level modeling, totaling 40 minutes of data per individual. The timing of the auditory stimulus was convolved with the canonical double-gamma hemodynamic response function to generate the auditory task regressor. The concatenated data were modeled using a general linear model (GLM) and processed using AFNI’s *3dREMLfit*. This ME-ICA model included six motion regressors and their derivatives, HRF-convolved end-tidal CO_2_, the auditory task regressor, the rejected ICA components and up to fourth-order polynomial terms.

#### 2.3.5 Statistical Analyses

Data were analyzed in each participant’s functional space using whole-brain and region-specific approaches. All subject-specific maps and activation results were then transformed to each subject’s anatomical space for visualization (*epi_reg, FLIRT,* FSL).

A schematic of the whole-brain and region-specific analyses used in this study can be found in Figure 1A.

**Figure 1:**
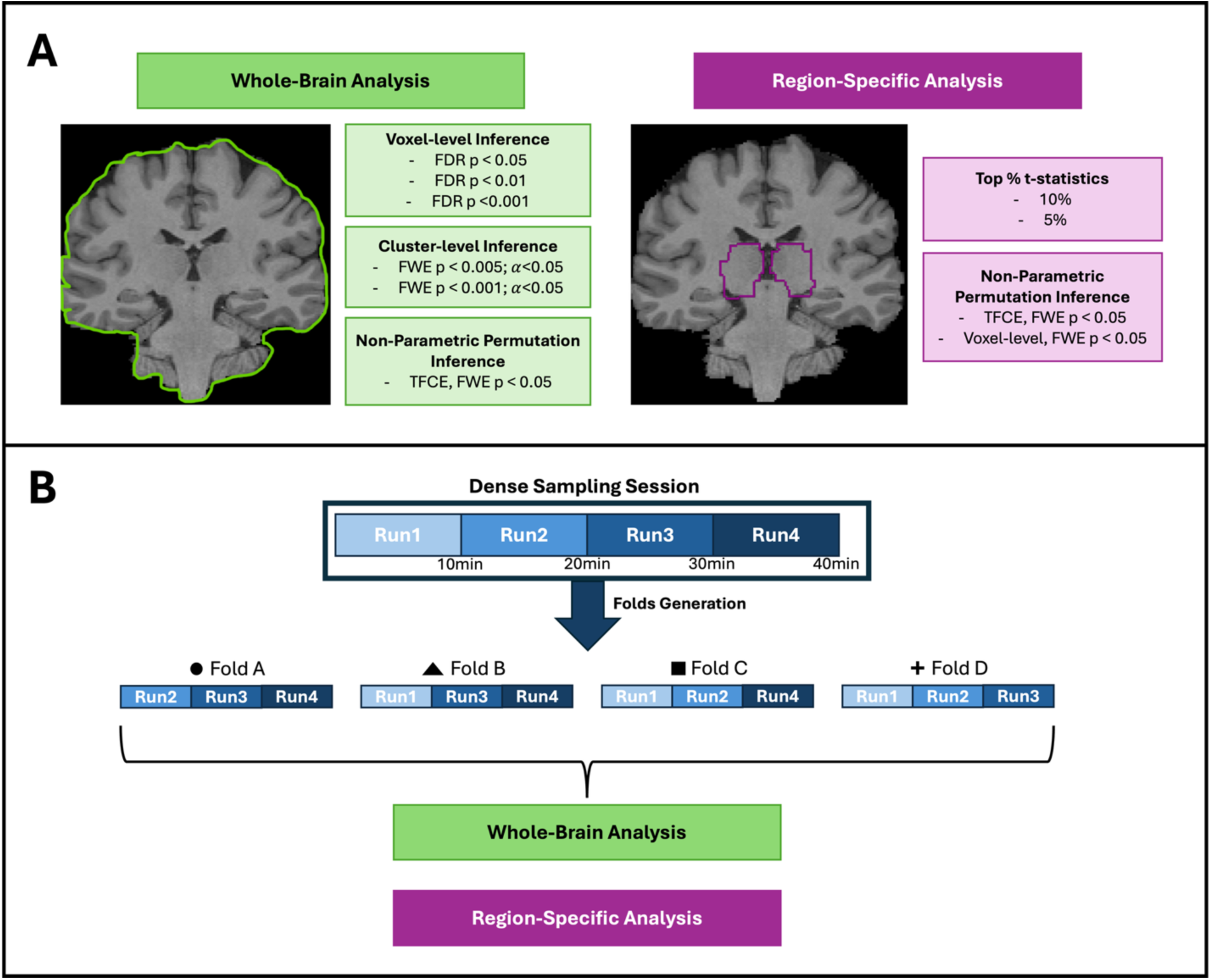
(A) Whole-brain and region-specific analyses for densely sampled single-subject datasets. Voxel-level inference testing employed a false discovery rate (FDR) correction using three different thresholds (*p* < 0.05, *p* < 0.01, and *p* < 0.001). Cluster-level inference testing performed Family-wise error (FWE) correction with clustering at *α* < 0.05 and two thresholds (*p* < 0.005 and *p* < 0.001). Non-parametric permutation inference employed threshold free cluster enhancement (TFCE) with Family-wise error (FWE) correction (*p* < 0.05). Region-specific analysis with top % t-statistics results were found by extracting the top 5% and 10% t-statistics within masks of thalamic, cerebellar, and brainstem regions. These masks were also employed for the non-parametric permutation approach with TFCE and voxel-level inference; both used a FWE correction (*p* < 0.05). **(B)** Schematic of the creation of the 30-minute folds. Folds were generated by concatenating three out of the four available functional runs. The concatenation order was varied, yielding four folds per subject. Each fold was analyzed using the approaches detailed in panel A.

##### 2.3.5.1 Whole-Brain Analyses

###### Voxel-Level Inference

Significant voxels of positive activation for the concatenated results (i.e. using all 40 minutes of scan data) were found in a whole-brain mask in functional space correcting for multiple comparisons using False Discovery Rate at three different thresholds (*p* < 0.05, *p* < 0.01, and *p* < 0.001), named VoxFDR_p05, VoxFDR_p01, VoxFDR_p001, respectively.

###### Cluster-Level Inference

Activation maps of the concatenated results were obtained by performing a whole-brain one-sided right-tail t-test (3dFWHMx, 3dClustSim, 3dClusterize), clustered at *α* < 0.05 and thresholded at both *p* < 0.005 (ClustFWE_p005) and *p* < 0.001 (ClustFWE_p001).

###### Non-parametric Permutation Inference

To perform non-parametric permutation on single-subject data, special care must be taken to avoid violating the required assumption for this type of inference testing: exchangeability (Lindquist & Mejia, 2015). To account for the repeated scans from each individual, each functional scan was first modeled independently using the GLM detailed in Section 2.3.4 and then each scan’s beta coefficient maps were combined using runMerge. This combined beta coefficient map was analyzed using FSL’s RANDOMISE. Permutations were restricted using exchangeability blocks detailing that all scans belonged to the same individual. The activation maps were then obtained by performing a one-sample t-test with threshold-free cluster enhancement (TFCE) and family-wise error correction (*p* < 0.05) within a whole brain mask in functional space.

##### 2.3.5.2 Region-Specific Analyses

Region-specific analyses have been previously conducted to better elucidate activation in challenging subcortical regions (Medina et al., 2026; Oliva et al., 2022; Reddy et al., 2024) and its benefits have been shown in the identification of activation of auditory nuclei (Medina et al., 2026). Thus, we employed this methodology to boost sensitivity in regions known *a priori* to be involved with auditory processing (i.e. thalamus, cerebellum, and brainstem) (E et al., 2012; Medina et al., 2026; Petacchi et al., 2005; Sigalovsky & Melcher, 2006; Yetkin et al., 2004). The thalamus and brainstem masks were created by thresholding these regions from the Harvard-Oxford Subcortical Structural Atlas at 50% and 25%, respectively. The cerebellar mask was created by thresholding the MNI structural atlas region at 50%. Region masks were then transformed to each subject’s functional space.

###### Top Percentage t-statistics

As previously done to better localize the areas of most dominant activation (Medina et al., 2026), voxels with the top 5% and 10% t-statistics within masks of the thalamus, cerebellum, and brainstem were extracted from the concatenated results maps and denoted as Top5%_tstats and Top10%_tstats, respectively.

###### Non-parametric Permutation Inference

To perform non-parametric permutation on single-subject data, special care must be taken to avoid violating the required assumption for this type of inference testing: exchangeability (Lindquist & Mejia, 2015). To account for the repeated scans from each individual, each functional scan was first modeled independently using the GLM detailed in Section 2.3.4 and then each scan’s beta coefficient maps were combined using runMerge. This combined beta coefficient map was analyzed using FSL’s RANDOMISE. Permutations were restricted using exchangeability blocks detailing that all scans belonged to the same individual. The activation maps were then obtained by performing a one-sample t-test with TFCE and voxel-based thresholding with family-wise error correction (*p* < 0.05) within masks of each region detailed above.

#### 2.3.6 Stability of significant activation with data subsets

As it has been previously shown that 30 minutes of concatenated ME-ICA data may be sufficient to obtain bilateral auditory activation in thalamic, cerebellar, and brainstem regions (Medina et al., 2026), data subsets (i.e. “folds”) were generated to assess the robustness and stability of each method. For each method, we count the number of folds without activation clusters to evaluate sensitivity in expected regions. Additionally, we use difference in center of mass (COM) and dice similarity coefficient (DSC) to compare the results using all available data (i.e. 40 minutes of concatenated data) and the data subsets for each method. For simplicity, the activation clusters found using the 40 minutes of data will be denoted as “AC_total” and those found using the 30-minute folds will be denoted as “AC_fold”. Given that non-parametric permutation approaches did not yield significant activation in whole-brain nor region-specific analyses, they were excluded from the subsequent analyses.

##### Creation of 30-minute Folds

Folds of 30-minute data were generated by concatenating three functional runs. The concatenation order was varied, ultimately yielding four 30-minute folds per individual – a schematic detailing each fold’s generation can be found on ***Figure 1B***. Each fold was modeled using the GLM described in Section 3.3.4 and analyzed using the approaches detailed in 3.3.5.

##### Number of Folds without Activation Clusters

The number of folds that did not yield any activation clusters were counted for each inference method. To account for the bilateral nature of the auditory nuclei, results were plotted for both Left and Right hemispheres within masks of the thalamus, region of cerebellar lobules VIIb/VIIIa, and midbrain. Folds corresponding to sub-04* appear as dashed areas to note the earbud malfunction for this participant. (See Results for more details on this participant.) To better localize the pertinent cerebellar activation, a mask for the cerebellar lobules VIIb/VIIIa was created using the MNI FNIRT atlas, thresholded at 50% and dilated by 2 voxels. The midbrain mask was created by manually eroding all voxels below the superior pontine sulcus from the thresholded brainstem mask detailed above. Ideally, a robust inference scheme would identify activation clusters on all folds (4 folds/subject; 20 folds total).

##### Difference in Center of Mass

To evaluate the stability of each method, differences in center of mass (COM) were found by calculating the Euclidean distance between the COM of AC_total and each AC_fold’s COM. The difference in COM was calculated for both Left and Right hemispheres within masks of the thalamus, region of cerebellar lobules VIIb/VIIIa, and midbrain. For each region and hemisphere, significant differences in the COM values between methods were tested as Difference in COM ∼ Method + (1|Subject); p-value was Bonferroni-corrected for multiple comparisons.

##### Dice Coefficient Scores

To assess spatial overlap between AC_total and each AC_fold, DSC were calculated within masks of the thalamus, cerebellum, and midbrain. For each region and hemisphere, significant differences in the DSC between methods were tested as DSC ∼ Method + (1|Subject); p-value was Bonferroni-corrected for multiple comparisons.

## 3. Results

All scans were successfully collected as described above. It must be noted that sub-04* experienced intermittent delivery of the auditory stimulus during scanning due to malfunction of one earbud and thus results from this participant must be interpreted with caution. An asterisk was added in each figure as a reminder of this note.

### 3.1 Cortical and Thalamic Activation with Whole-Brain Analyses

Across all subjects, only VoxFDR_p05 identified significant bilateral activation in expected cortical and subcortical auditory regions - primary auditory cortex (PAC) and medial geniculate nuclei (MGN), respectively (***Figure 2***). For subjects with prominent auditory activity (i.e. large activation extent in MGN), such as sub-03 and sub-05, both voxel-level and cluster-level inference testing identified significant bilateral activation in PAC and MGN, regardless of significance thresholding. The non-parametric permutation inference approach did not identify any significant activation, even in large cortical areas like PAC, suggesting that 4 runs (10 min/run) may not be sufficient data to power this type of analysis for densely sampled single-subject inference. Subthreshold (i.e., not statistically significant) activation is present in PAC for all subjects and in MGN for three subjects (sub-03, sub-04*, and sub-05).

**Figure 2:**
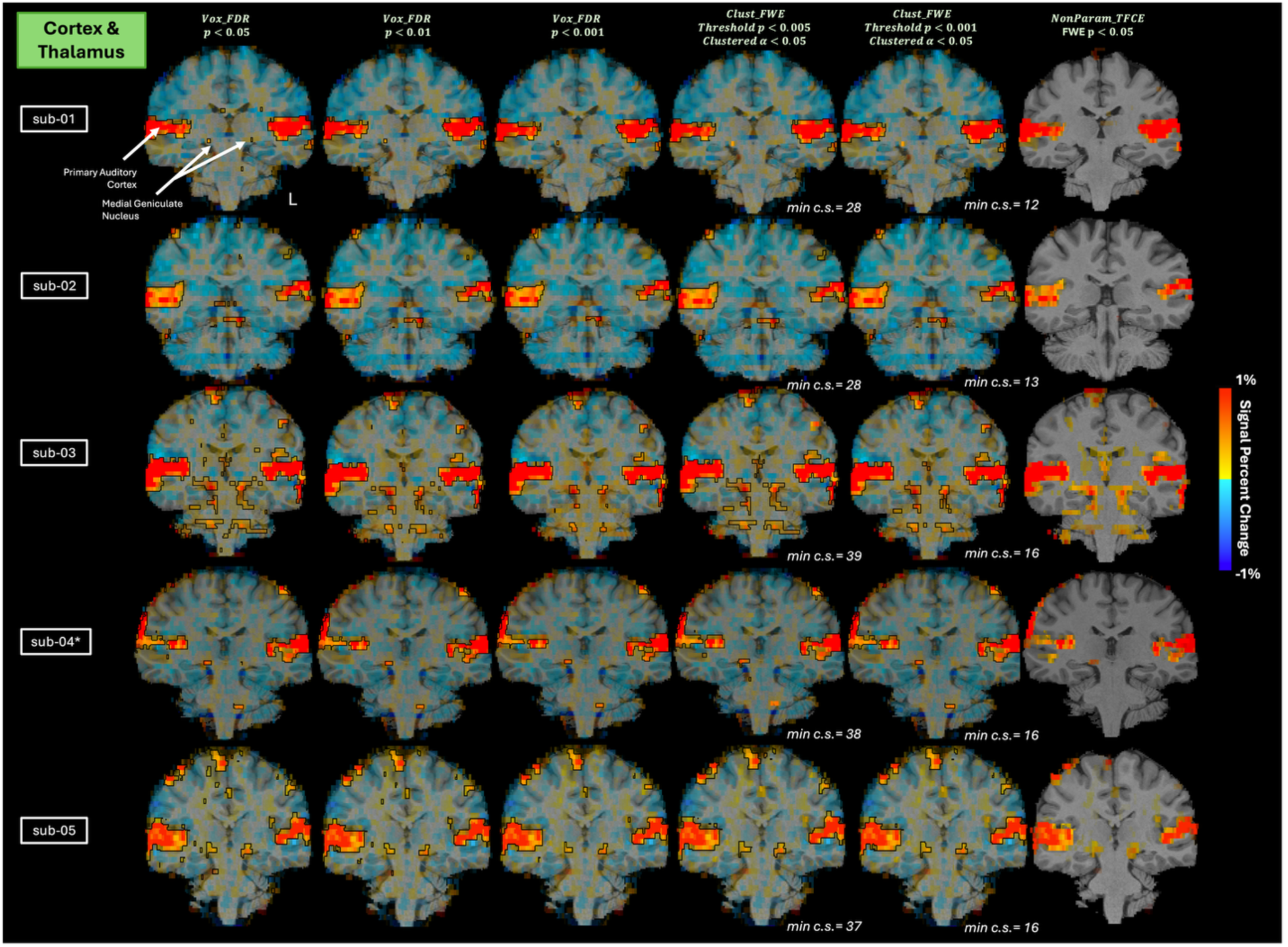
Whole-brain analyses results for cortex and thalamus using voxel-level inference (FDR with *p* < 0.05, *p* < 0.01, and *p* < 0.001), cluster-level inference (one-sided right-tail t-test, FWE thresholding at *p* < 0.005 and *p* < 0.001, and clustering at *α* < 0.05), and non-parametric methods (RANDOMISE, one-sided t-test, TFCE, FWE *p* < 0.05). The opacity of the beta-coefficient values was modulated by t-statistic. Black outlines showcase significant clusters of positive activation, compared to rest. The minimum cluster size (min c.s.) is shown for each subject.

Overall, although more conservative significance thresholding may remove spurious clusters of activation, it may additionally remove expected activation in subcortical regions. The conventional voxel-level inference scheme with a less conservative threshold (i.e. FDR *p* < 0.05) might be more suitable when investigating regions with small nuclei (Glendenning & Masterton, 1998; Sitek et al., 2019) and poor contrast (De Martino et al., 2015; Saranathan et al., 2021; Williams et al., 2024) such as the thalamus.

### 3.2 Thalamic Activation with Region-Specific Analyses

For all subjects except sub-04* (who reported earphone issues after the scanning session was completed), top5%_tstats and top10%_tstats identified similar bilateral activation in MGN (***Figure 3***). Additionally, the activation clusters for both top5%_tstats and top10%_tstats were located in the same slices and were of comparable size to the whole-brain results in ***Figure 2***. Non-parametric permutation methods did not identify significant activation for any subject. Subthreshold bilateral activation is present in MGN areas for three subjects (sub-03, sub-04*, and sub-05) for both TFCE and voxel-level inference methods.

**Figure 3:**
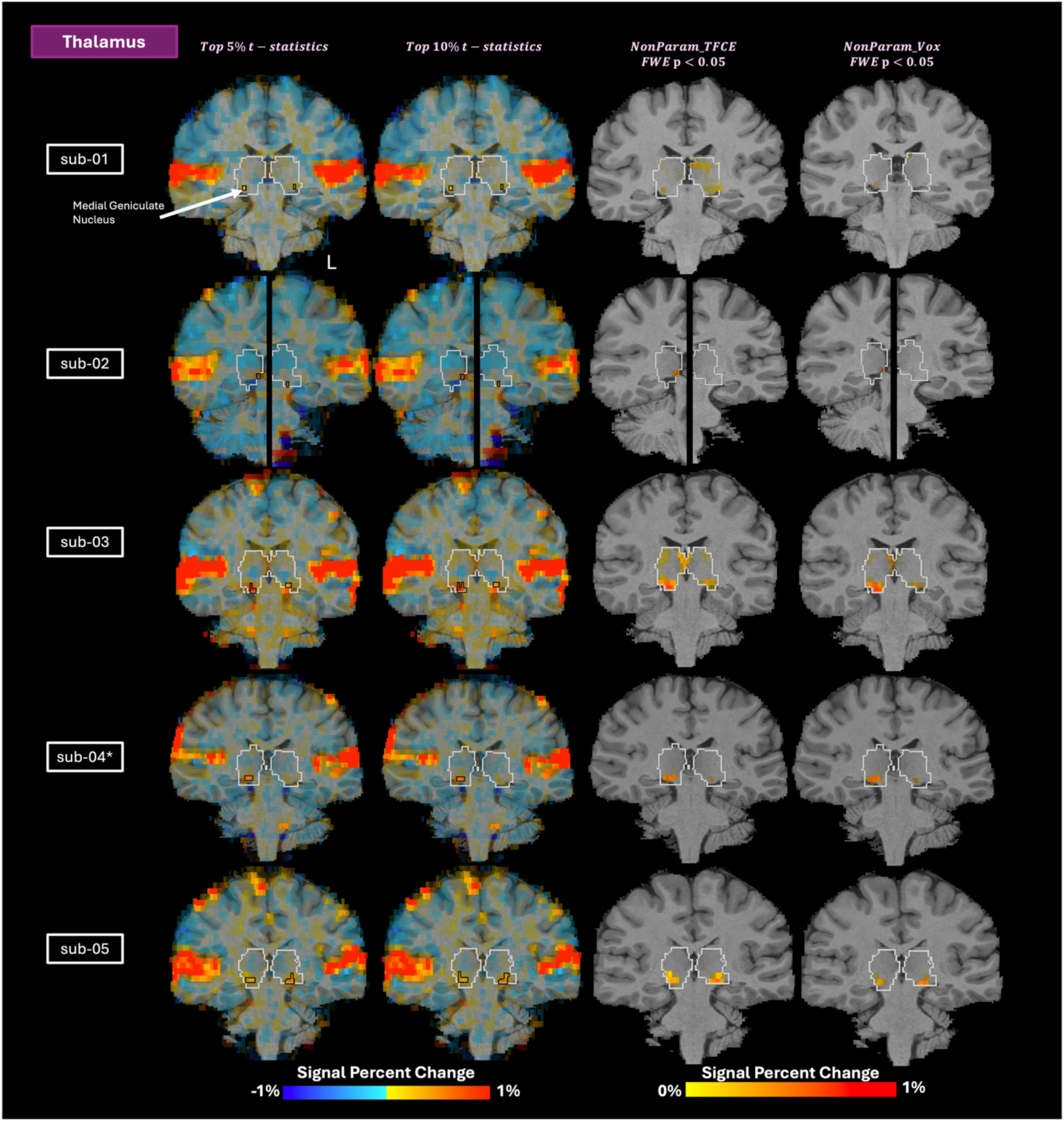
Region-specific analyses results for the thalamus using top percentage of t-statistics (5% and 10%, uncorrected) and non-parametric methods (RANDOMISE, one-sided t-test, FWE *p* < 0.05 with TFCE and voxel-level inference). The opacity of the beta-coefficient values was modulated by t-statistic. Black outlines showcase clusters of activation. Silver outlines denote the thalamic regions used for each analysis.

### 3.3 Cerebellar Activation with Whole-Brain Analyses

Regardless of significance thresholding, voxel-based and cluster-based inference methods identified bilateral cerebellar activation in lobules VIIb/VIIIa for all subjects except sub-02 with ClustFWE_p005, as shown in ***Figure 4***. This finding suggests that the cerebellum might yield more robust activation than other subcortical regions (e.g. thalamus), reaching significance even with more conservative thresholding and different inference schemes likely due to the larger size of these lobules (Lyu et al., 2024). The non-parametric method did not identify significant activation for any subject. Subthreshold bilateral activation is present in lobules VIIb/VIIIa areas for four subjects (sub-01, sub-03, sub-04*, and sub-05) for both TFCE and voxel-level inference methods.

**Figure 4:**
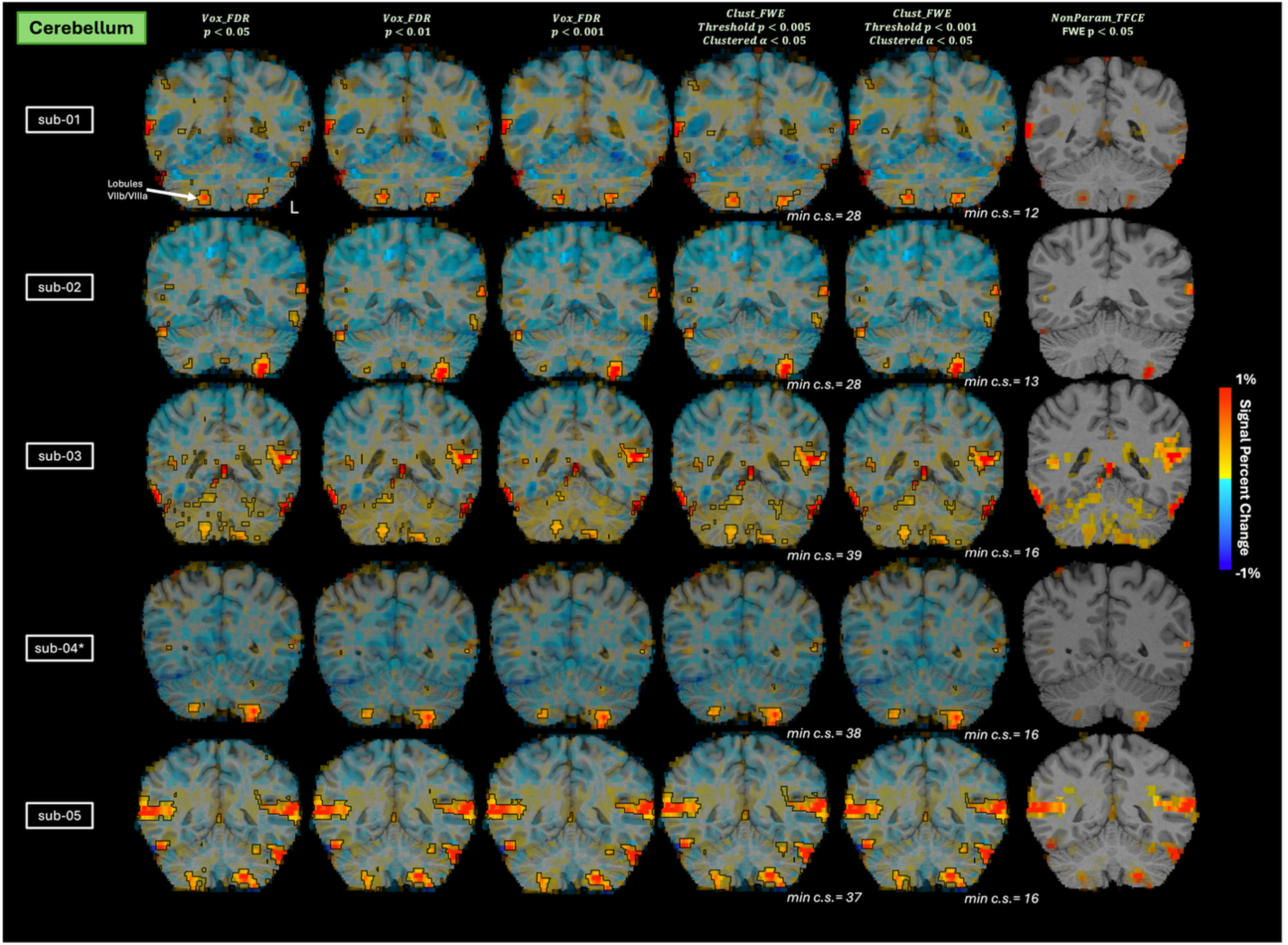
Whole-brain analyses results for the cerebellum using voxel-level inference (FDR with *p* < 0.05, *p* < 0.01, and *p* < 0.001), cluster-level inference (one-sided right-tail t-test, FWE thresholding at *p* < 0.005 and *p* < 0.001, and clustering at *α* < 0.05), and non-parametric methods (RANDOMISE, one-sided t-test, TFCE, FWE *p* < 0.05). The opacity of the beta-coefficient values was modulated by t-statistic. Black outlines showcase significant clusters of positive activation, compared to rest. The minimum cluster size (min c.s.) is shown for each subject.

### 3.4 Cerebellar Activation with Region-Specific Analyses

For all subjects except sub-02, top5%_tstats and top10%_tstats identified bilateral activation in cerebellar lobules VIIb/VIIIa (***Figure 5***). Non-parametric methods did not identify significant activation for any subject. Subthreshold bilateral activation is present in lobules VIIb/VIIIa for four subjects (sub-01, sub-03, sub-04*, and sub-05) for both TFCE and voxel-level inference methods. For sub-03, the TFCE approach yielded spurious clusters of subthreshold activation in areas not expected to be involved with auditory processing. This finding suggests that NonParam_TFCE may fail to appropriately delineate primary regions of task-based activation for individuals with widespread activation, such as sub-03, possibly due to this method’s increased sensitivity and spatial bias (Noble et al., 2020).

**Figure 5:**
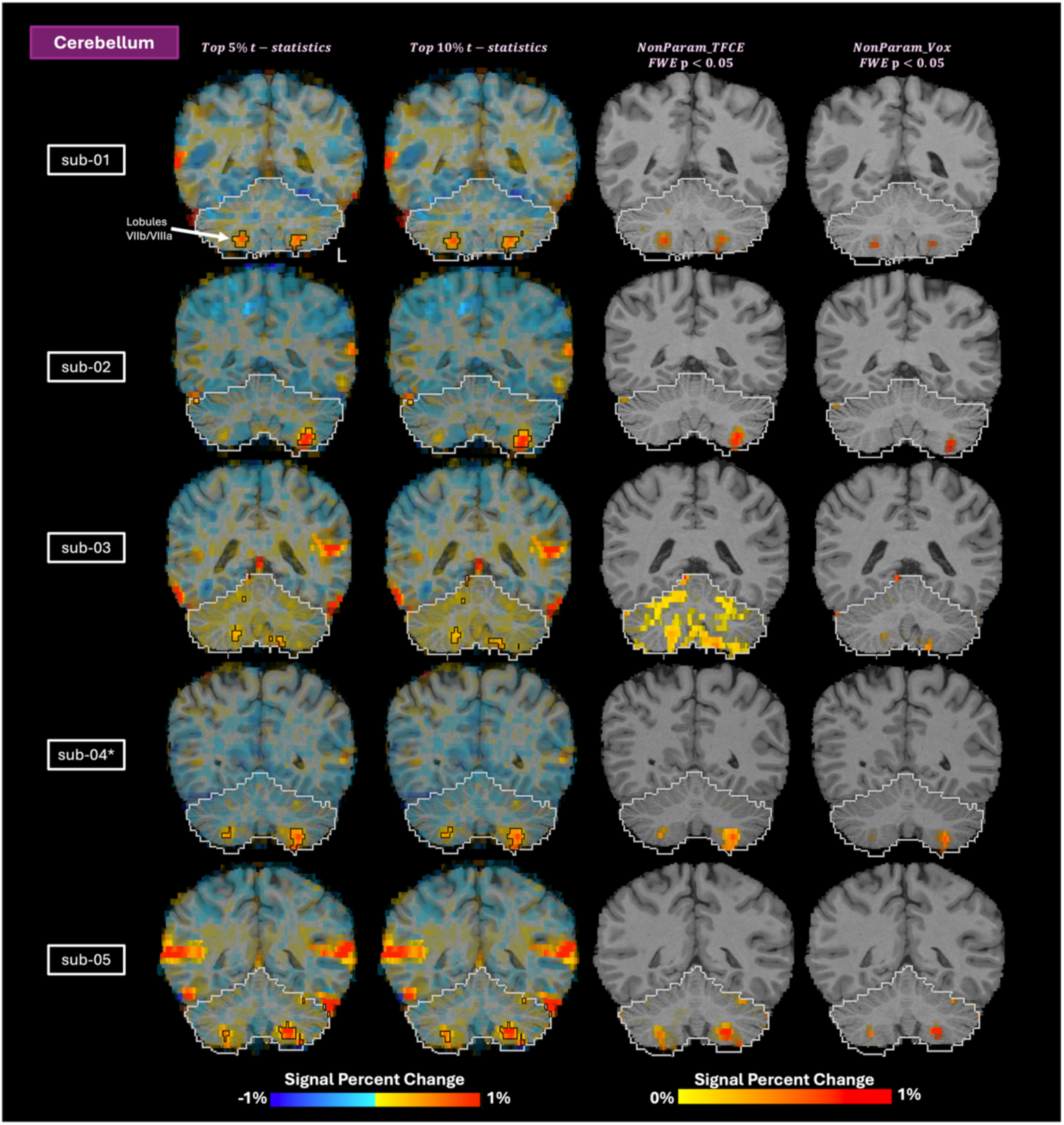
Region-specific analyses results for the cerbellum using top percentage of t-statistics (5% and 10%, uncorrected) and non-parametric methods (RANDOMISE, one-sided t-test, FWE *p* < 0.05 with TFCE and voxel-level inference). The opacity of the beta-coefficient values was modulated by t-statistic. Black outlines showcase clusters of activation. Silver outlines denote the cerebellar region used for each analysis.

### 3.5 Brainstem Activation with Whole-Brain Analyses

For all subjects except sub-04* (who reported earphone issues), VoxFDR_p05 identified significant bilateral activation in the inferior colliculi (IC), as seen on ***Figure 6***. Three subjects (sub-02, sub-03, and sub-05) exhibited significant bilateral activation in IC for both voxel-level and cluster-level inference methods, regardless of significance thresholding, with the exception of sub-05 with VoxFDR_p001. It is important to note that all voxel-level and cluster-level inference methods identified significant unilateral activation in IC for sub-04*, whose earbud malfunctioned during data collections, suggesting that this might be the reason behind the lack of bilateral activation. The non-parametric method did not identify any significant activation. Subthreshold bilateral activation is present in IC for two subjects (sub-03 and sub-05).

**Figure 6:**
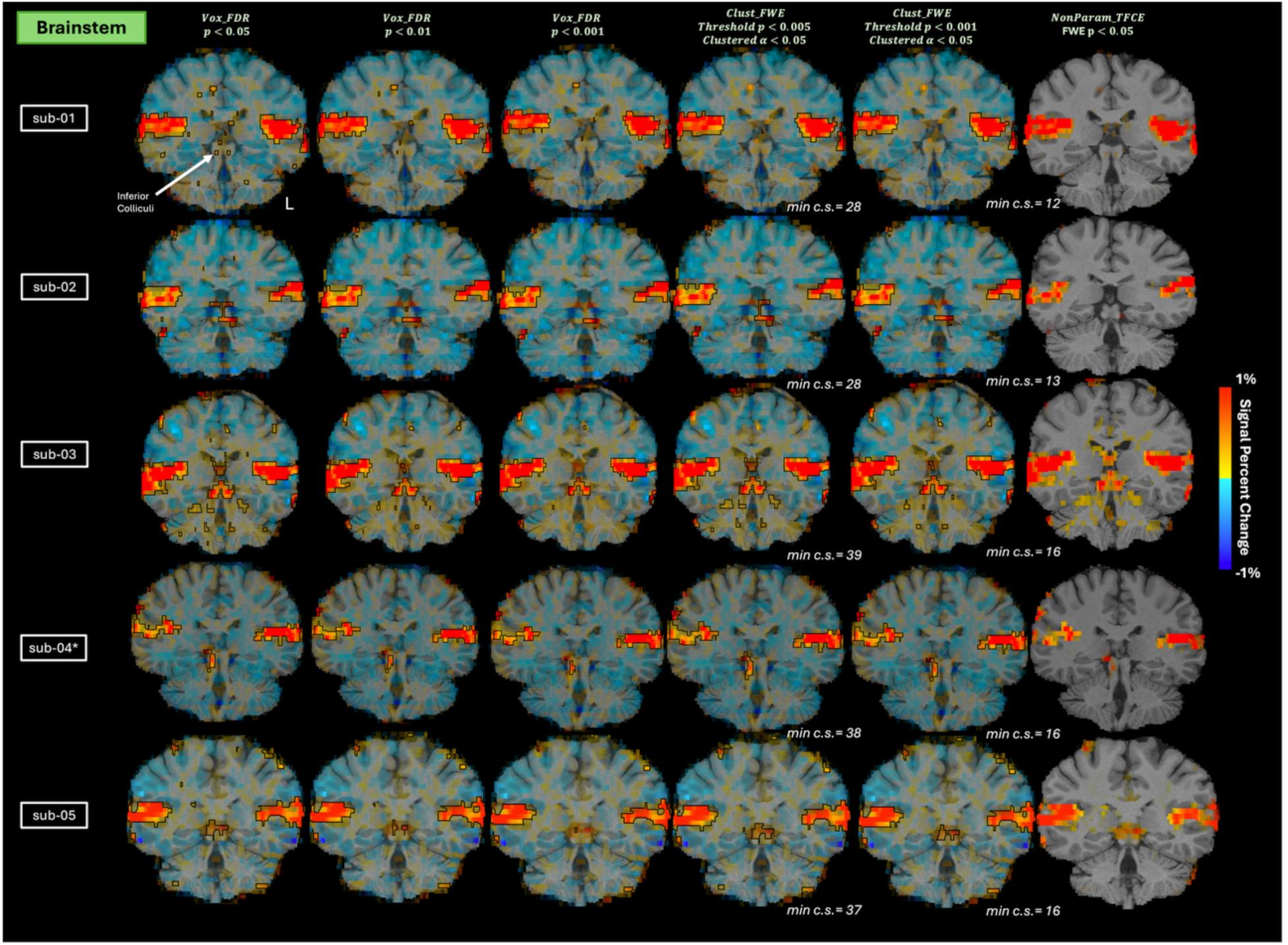
Whole-brain analyses results for the brainstem using voxel-level inference (FDR with *p* < 0.05, *p* < 0.01, and *p* < 0.001), cluster-level inference (one-sided right-tail t-test, FWE thresholding at *p* < 0.005 and *p* < 0.001, and clustering at *α* < 0.05), and non-parametric methods (RANDOMISE, one-sided t-test, TFCE, FWE *p* < 0.05). The opacity of the beta-coefficient values was modulated by t-statistic. Black outlines showcase significant clusters of positive activation, compared to rest. The minimum cluster size (min c.s.) is shown for each subject.

### 3.6 Brainstem Activation with Region-Specific Analyses

For all subjects, except sub-04*, top5%_tstats and top10%_tstats identified bilateral activation in IC (***Figure 7***). Non-parametric permutation methods did not identify significant activation for any subject. Subthreshold bilateral activation is present in IC area for just one subject (sub-03) for both TFCE and voxel-level inference methods.

**Figure 7:**
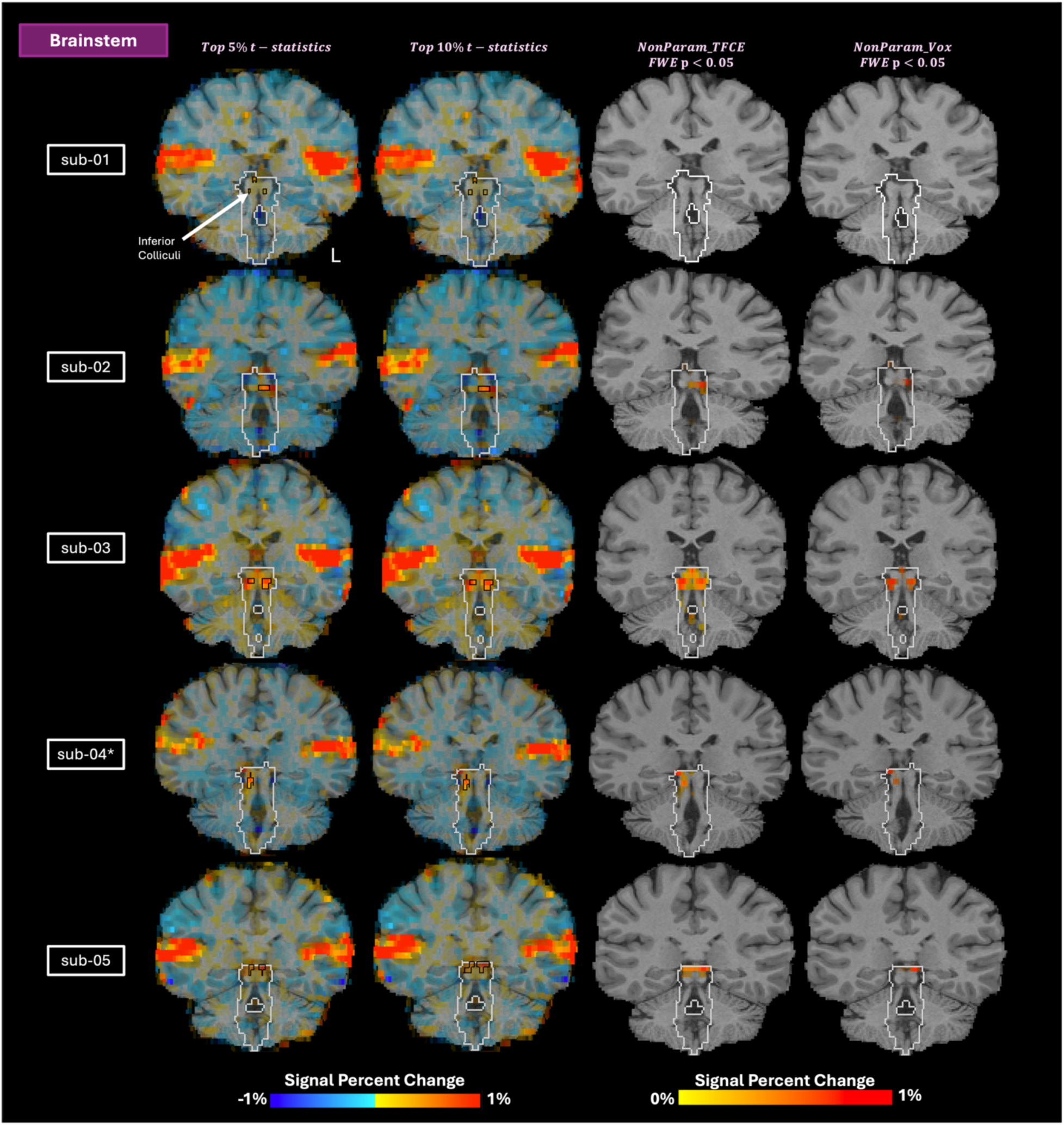
Region-specific analyses results for the brainstem using top percentage of t-statistics (5% and 10%, uncorrected) and non-parametric methods (RANDOMISE, one-sided t-test, FWE *p* < 0.05 with TFCE and voxel-level inference). The opacity of the beta-coefficient values was modulated by t-statistic. Black outlines showcase clusters of activation. Silver outlines denote the thalamic regions used for each analysis.

### 3.7 Evaluation of Stability across Subsets of the Densely Sampled Data

Across regions, VoxFDR_p05 yielded the least number of folds with no activation clusters, suggesting that this approach might provide more robust identification of small auditory nuclei, even with 30-minutes of functional data (***Figure 8A***). Average difference in COM and DSC values can be found in ***Table 1*** and ***Table 2***, respectively. Results were plotted in ***Figure 8B*** and ***Figure 8C***, respectively. Ideally, a reliable approach would yield a low difference in COM and a high DSC – see ***Supplemental Figure 1*** for example cases. In the thalamus, ClustFW_p001 yielded the smallest mean Euclidean distance (2.0 mm) between COMs and highest mean DSC (0.78) across both hemispheres. In the cerebellum, the top t-statistic approaches yielded the smallest mean Euclidean distance (0.84 mm; Top5%_tstat) between COMs and highest DSC (0.85; Top10%_tstat) across both hemispheres. In the midbrain, VoxFDR_p05 yielded the smallest mean Euclidean distance (1.1 mm) between COMs and highest mean DSC (0.74) across both hemispheres.

**Figure 8:**
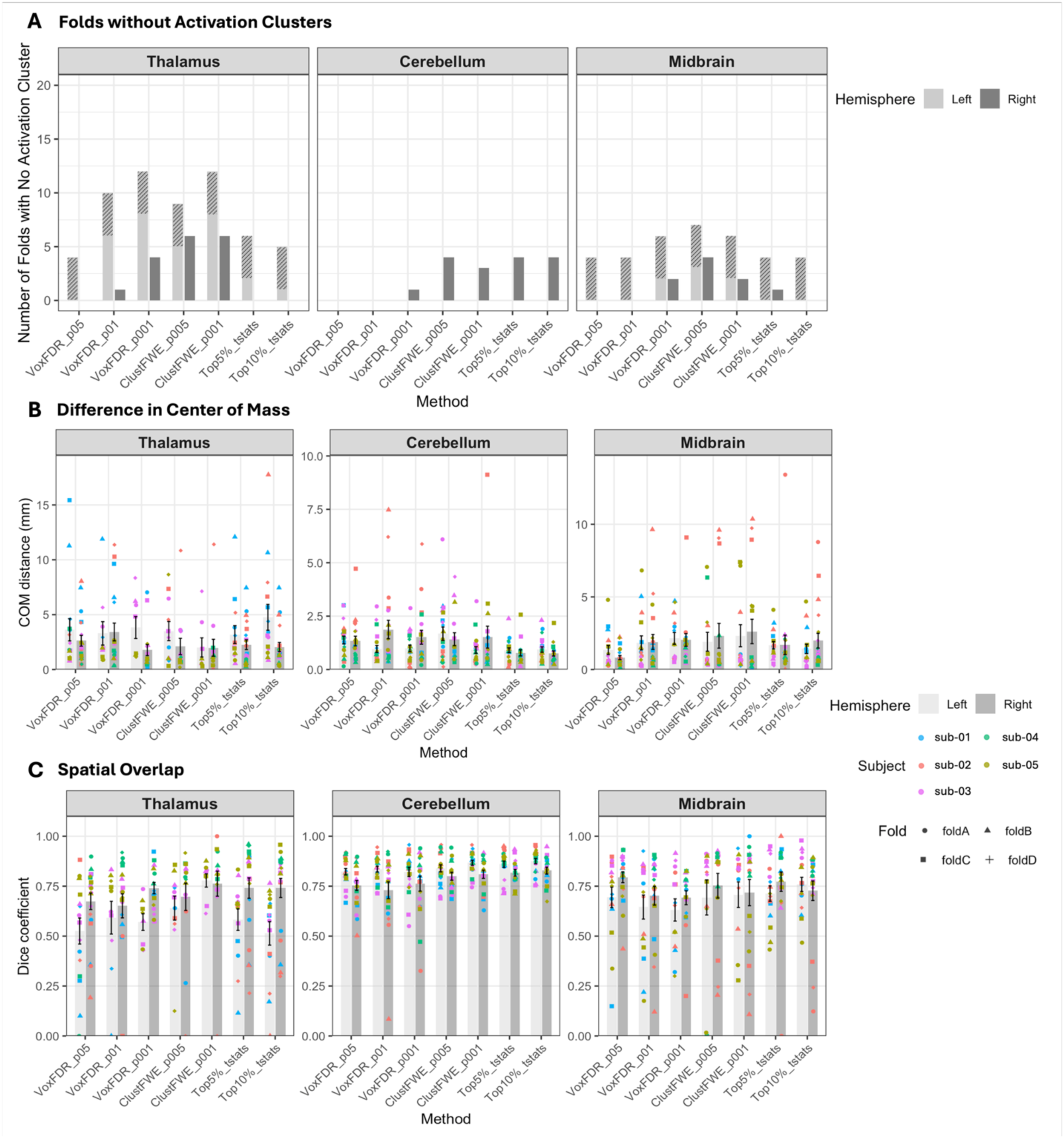
(A) Number of folds with no activation clusters in Left and Right hemispheres of thalamic, cerebellar, and midbrain regions. Folds of 30-minutes of data were generated for each subject by concatenating three out of the four available functional runs. The concatenation order was varied yielding four folds for each subject (i.e. 20 folds total). Folds corresponding to sub-04* appear as dashed areas to note the earbud malfunction for this participant. Activation clusters found using all 40-minutes of densely sampled data were denoted as “AC_total” and those found using the 30-minute folds were denoted as “AC_fold”. **(B)** Difference in center of mass (COM) between AC_total and AC_ fold in thalamic (MGN), cerebellar (lobules VIIb/VIIIa), and midbrain (IC) regions. The Euclidian distance between the COMs for Left and Right hemispheres are plotted, color-coded based on subject and shape is determined by fold. **(C)** Spatial overlap between AC_total and AC_ fold in thalamic (MGN), cerebellar (lobules VIIb/VIIIa), and midbrain (IC) regions. The dice coefficient score for Left and Right hemispheres are plotted, color-coded based on subject and shape is determined by fold.

**Table 1:**
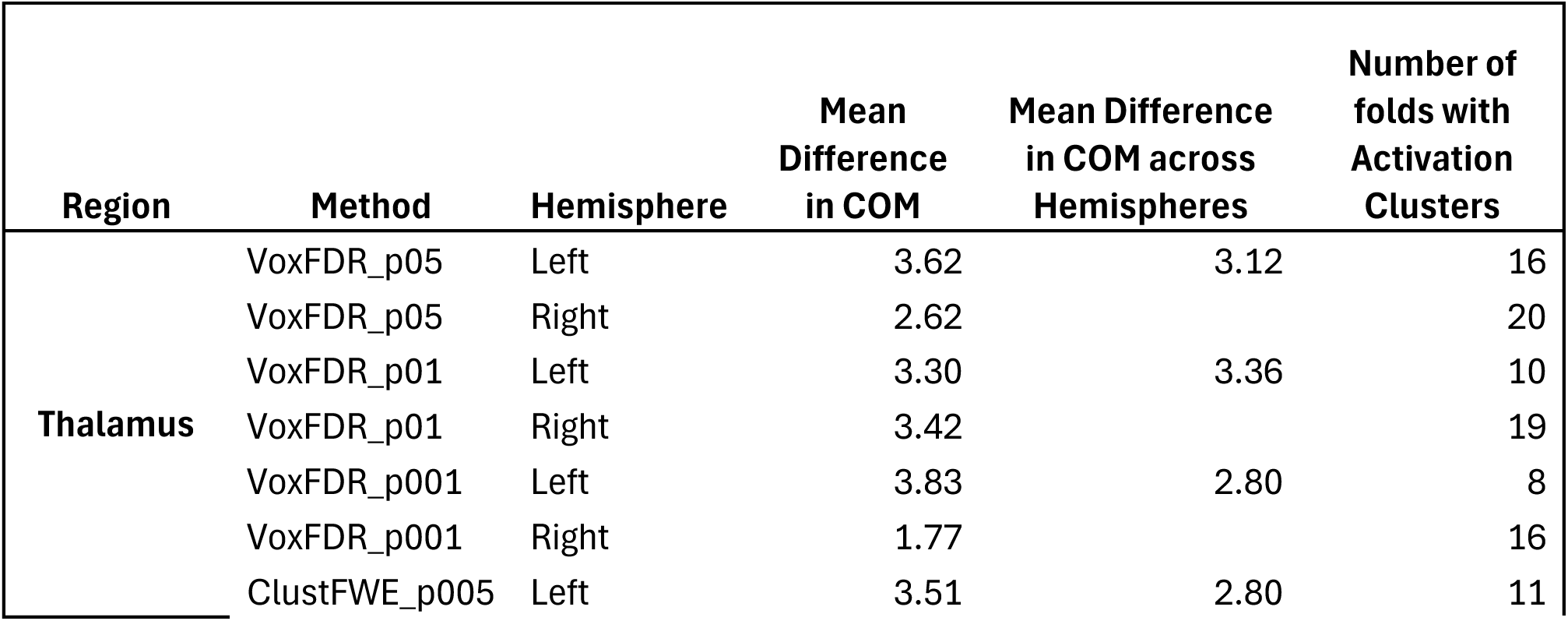
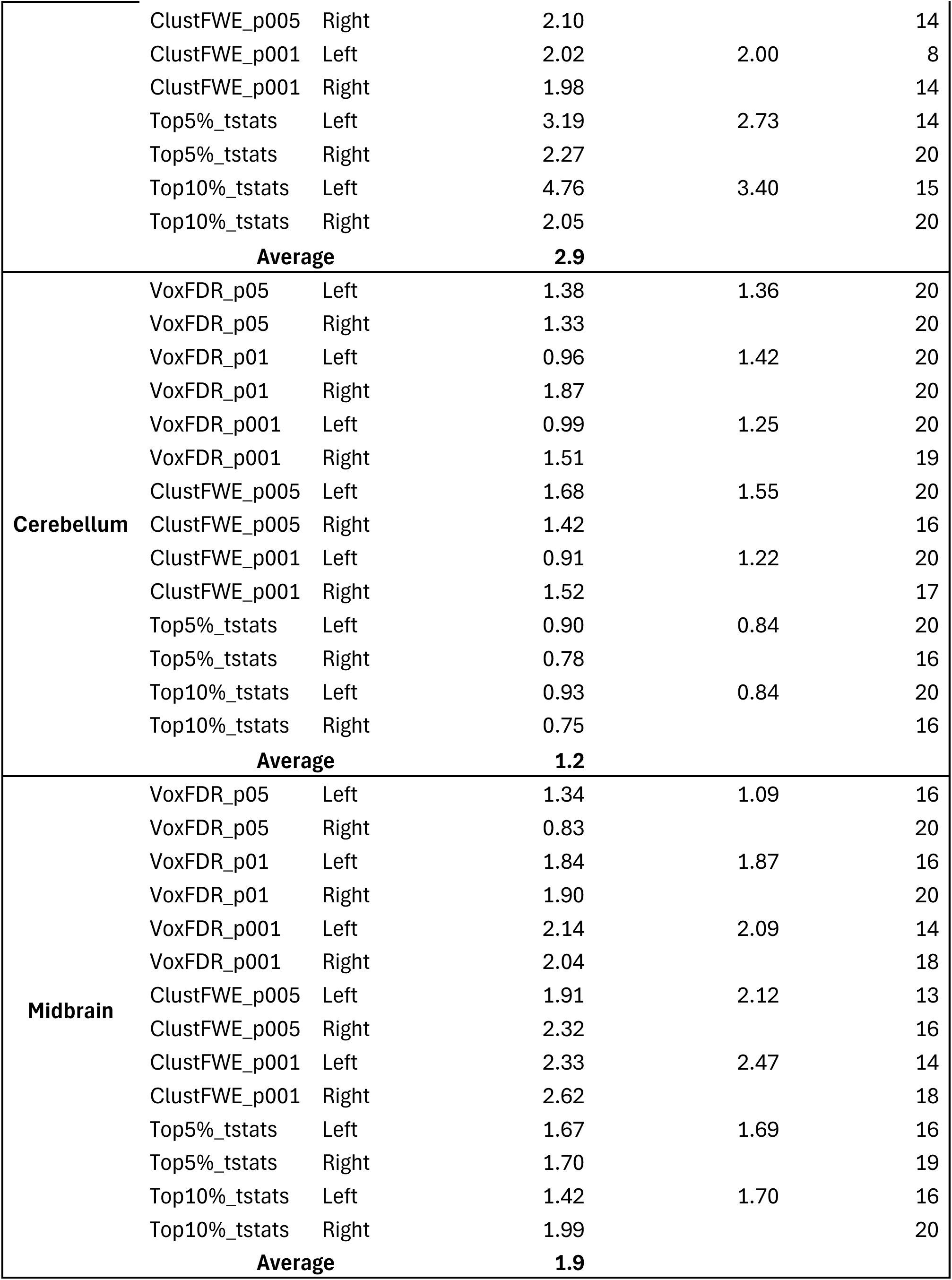
Mean difference in center of mass (COM) between AC_total and all AC_ folds in thalamic (MGN), cerebellar (lobules VIIb/VIIIa), and midbrain (IC) regions. The Euclidian distance between the COMs for Left, Right and Both hemispheres are reported. Total number of folds with activation clusters is shown for each hemisphere. Only folds with activation clusters were used in all calculations.

**Table 2:** Mean dice similarity coefficient (DSC) between AC_total and all AC_ folds in thalamic (MGN), cerebellar (lobules VIIb/VIIIa), and midbrain (IC) regions. Mean DSC values for Left, Right and Both hemispheres are reported. Total number of folds with activation clusters is shown for each hemisphere. Only folds with activation clusters were used in all calculations.

| Region | Method | Hemisphere | Mean DSC | Mean DSC across Hemispheres | Number of folds with Activation Clusters |
| --- | --- | --- | --- | --- | --- |
| Thalamus | VoxFDR_p05 | Left | 0.53 | 0.60 | 16 |
|  | VoxFDR_p05 | Right | 0.67 |  | 20 |
|  | VoxFDR_p01 | Left | 0.59 | 0.62 | 10 |
|  | VoxFDR_p01 | Right | 0.65 |  | 19 |
|  | VoxFDR_p001 | Left | 0.57 | 0.66 | 8 |
|  | VoxFDR_p001 | Right | 0.74 |  | 16 |
|  | ClustFWE_p005 | Left | 0.64 | 0.67 | 11 |
|  | ClustFWE_p005 | Right | 0.70 |  | 14 |
|  | ClustFWE_p001 | Left | 0.78 | 0.77 | 8 |
|  | ClustFWE_p001 | Right | 0.76 |  | 14 |
|  | Top5%_tstats | Left | 0.58 | 0.66 | 14 |
|  | Top5%_tstats | Right | 0.74 |  | 20 |
|  | Top10%_tstats | Left | 0.51 | 0.63 | 15 |
|  | Top10%_tstats | Right | 0.74 |  | 20 |
|  | Average |  | 0.66 |  |  |
| Cerebellum | VoxFDR_p05 | Left | 0.82 | 0.79 | 20 |
|  | VoxFDR_p05 | Right | 0.75 |  | 20 |
|  | VoxFDR_p01 | Left | 0.84 | 0.78 | 20 |
|  | VoxFDR_p01 | Right | 0.73 |  | 20 |
|  | VoxFDR_p001 | Left | 0.82 | 0.79 | 20 |
|  | VoxFDR_p001 | Right | 0.76 |  | 19 |
|  | ClustFWE_p005 | Left | 0.84 | 0.82 | 20 |
|  | ClustFWE_p005 | Right | 0.80 |  | 16 |
|  | ClustFWE_p001 | Left | 0.87 | 0.84 | 20 |
|  | ClustFWE_p001 | Right | 0.81 |  | 17 |
|  | Top5%_tstats | Left | 0.86 | 0.84 | 20 |
|  | Top5%_tstats | Right | 0.82 |  | 16 |
|  | Top10%_tstats | Left | 0.88 | 0.85 | 20 |
|  | Top10%_tstats | Right | 0.83 |  | 16 |
| **Average** |  |  | **0.82** |  |  |
| **Midbrain** | VoxFDR_p05 | Left | 0.69 | 0.74 | 16 |
|  | VoxFDR_p05 | Right | 0.79 |  | 20 |
|  | VoxFDR_p01 | Left | 0.65 | 0.67 | 16 |
|  | VoxFDR_p01 | Right | 0.70 |  | 20 |
|  | VoxFDR_p001 | Left | 0.63 | 0.66 | 14 |
|  | VoxFDR_p001 | Right | 0.69 |  | 18 |
|  | ClustFWE_p005 | Left | 0.69 | 0.72 | 13 |
|  | ClustFWE_p005 | Right | 0.75 |  | 16 |
|  | ClustFWE_p001 | Left | 0.71 | 0.71 | 14 |
|  | ClustFWE_p001 | Right | 0.72 |  | 18 |
|  | Top5%_tstats | Left | 0.71 | 0.74 | 16 |
|  | Top5%_tstats | Right | 0.77 |  | 19 |
|  | Top10%_tstats | Left | 0.76 | 0.74 | 16 |
|  | Top10%_tstats | Right | 0.73 |  | 20 |
| **Average** |  |  | **0.71** |  |  |

Across regions, the difference in COM and DSC values between folds were averaged for each hemisphere and method. For all methods, this mean difference in COM was 2.9, 1.2, and 1.9 mm for the thalamus, cerebellum, and midbrain, respectively. The mean DSC was 0.66, 0.82, and 0.71 for thalamus, cerebellum, and midbrain, respectively. Across regions, no significant differences were found between methods for either metric.

## 4. Discussion

While task-based precision mapping has tremendous potential in identifying participant-specific loci of functional processing, the field has not yet settled on an optimal approach for localizing meaningful systems-level fMRI responses. Thus, in this study, we evaluated whole-brain and region-specific inference testing approaches with varying statistical thresholds to investigate the capacity of each to identify task-based BOLD fMRI activation in expected regions of the cortical and subcortical auditory system and to characterize significance in densely sampled individual-subject data. Compared to all tested approaches, we found that a whole-brain voxel-level inference approach with FDR *p* < 0.05 was more sensitive at identifying significant bilateral activation across regions and may provide a more robust localization of small auditory nuclei, even with only 30-minutes of functional data. Furthermore, our results show that although qualitative region-specific approaches, such as top % t-statistics, do not test for traditional statistical significance, they may yield more consistent activation clusters with varying data length, especially in thalamic and cerebellar regions.

### 4.1 Cortical Activation

Across subjects, all whole-brain approaches yielded significant bilateral activation of PAC with the exception of the non-parametric permutation inference method. Although non-parametric methods have become a promising approach to characterize functional activity at the group-level (Medina et al., 2026; Oliva et al., 2022; Reddy et al., 2024), it is not widely used at the individual level as it requires additional considerations (Lindquist & Mejia, 2015), such as the use of exchangeability blocks to account for repeated scans. Our results show that this procedure might be underpowered with just 4 functional runs, yielding overly conservative statistics that prevent the identification of significant activation, even in large cortical clusters. This result is consistent across all regions for both whole-brain and region-specific analyses, implying that a non-parametric permutation approach may require greater amounts of individual-subject data for activation to reach significance.

### 4.2 Thalamic Activation

The thalamus is a region well-known for its poor anatomical contrast (De Martino et al., 2015; Saranathan et al., 2021; Williams et al., 2024) and small nuclei, such as the MGN (4x4x4 mm^3^) (Glendenning & Masterton, 1998; Sitek et al., 2019). Our results show that both a whole-brain analysis with voxel-level inference and region-specific top % of t-statistics analysis may appropriately overcome these challenges. Although the region-specific top % t-statistic approach does not test for traditional statistical significance, both 5% and 10% thresholds yielded comparable MGN activation clusters to the whole-brain voxel-level approach for all subjects, suggesting that this method might be an efficient exploratory approach to capture individual-specific dominant subcortical activation. Additionally, it is important to note that more conservative thresholds for the whole-brain voxel-level inference approach may remove bilateral clusters of activation, and thus a standard p<0.05 threshold might provide a more favorable sensitivity-specificity tradeoff for the detection of subcortical nuclei.

Although cluster-level inference methods yielded significant MGN clusters for some subjects, this approach may not be suitable for individuals with a smaller region of MGN activation as the minimum cluster size may be too large of a threshold for the identification of their subcortical nuclei. Despite evidence that cluster-level approaches may yield lower spatial specificity with large clusters (Lindquist & Mejia, 2015; Nichols, 2012; Woo et al., 2014), the three subjects who presented MGN activation showcased small clusters within the confines of the expected thalamic auditory nuclei, suggesting that reasonable specificity (Woo et al., 2014) might be provided by this approach in the thalamus when using densely sampled, single-subject data.

### 4.3 Cerebellar Activation

Recent studies have shown the involvement of cerebellar lobules VIIb/VIIIa during music tasks (E et al., 2012; Medina et al., 2026). Overall, our results show that robust activation is present in the cerebellum and significance may be reached even with more conservative thresholds, possibly due to the fact that these lobules extend a larger surface area (∼50x50mm^2^) (Lyu et al., 2024) compared to that of thalamic and brainstem auditory nuclei (4x4x4 mm^3^) (Glendenning & Masterton, 1998; Sitek et al., 2019). Bilateral activation of these lobules was identified by most approaches, with the exception of ClustFWE_p005, Top5%_tstats, and Top10%_tstats for sub-02, which only identified left lobule activity. It is worth emphasizing that left lateralized activity has been shown during auditory tasks (McLachlan & Wilson, 2017; Petacchi et al., 2005), especially with the left lobule VIII during music stimuli (E et al., 2012). Thus, this predominance in laterality might be a possible driver for this individual, especially with the Top5%_tstats and Top10%_tstats approaches which yield the most dominant activation in the inspected region.

The ability of the top % t-statistics method to tease out more focal and predominant cerebellar activation may be particularly useful when trying to define more specific regions of activity and remove spurious clusters of activation, especially for investigations with *a priori* knowledge of expected, functionally involved regions. As evidenced by sub-03 in ***Figure 4***, cerebellar results for whole-brain approaches may identify additional regions of activation which may not be task-related, and thus very stringent thresholds are necessary to detect the expected auditory activation. The challenge with using more stringent thresholds in whole-brain approaches is in the impact it may cause on other regions, such as the thalamus, where sensitivity of smaller nuclei is severely affected by uniformly applied conservative thresholds. Therefore, the top % t-statistic approach may be a complementary method to whole-brain approaches with thresholds that require additional balance when inspecting entire systems.

### 4.4 Brainstem Activation

The brainstem is a particularly challenging region given the presence of physiological and vascular confounds (Beissner, 2015). In addition, similar to thalamic auditory nuclei, brainstem auditory nuclei may be small in size (4x4x4 mm^3^) (Glendenning & Masterton, 1998; Sitek et al., 2019), and for certain subjects (e.g. sub-01), conservative thresholds and cluster-based approaches might fail to detect functional activity in this region. Nonetheless, despite these challenges, most whole-brain approaches (i.e. VoxFDR_p05, VoxFDR_p01, ClustFWE_p005, and ClustFWE_p001) identified significant bilateral IC activation for three out of the five subjects (i.e. sub-02, sub-03 and sub-05). Although activation cannot be classified as significant, a region-specific top % t-statistics approach, as well as other qualitative thresholding methods, have been successfully used in previous work (Medina et al., 2026; Salvo, Anderson, et al., 2025; Salvo, Lakshman, et al., 2025). Results show that both top % t-statistics thresholds identified bilateral IC activation for all subjects except for sub-04*, which presented unilateral IC activation. These findings suggest that a dense-sampling strategy might yield robust results even in this challenging region.

### 4.5 Evaluation of Stability across Subsets of the Densely Sampled Data

Similar to previous research (Medina et al., 2026), we found that the folds of 30-minute data successfully identified bilateral activation in thalamic, cerebellar, and brainstem auditory regions for most approaches and subjects. Across regions, the VoxFDR_p05 approach yielded the highest consistency, as evidenced by the least number of folds with no activation clusters (i.e. 4 folds in thalamus and midbrain). Notably, these folds belonged to sub-04*, who experienced an earbud malfunction during their scanning session. These results suggest that a whole-brain VoxFDR_p05 analysis might provide more robust identification of significant bilateral activation across different regions with lower scan duration.

To evaluate the stability of each method’s activation clusters, distance between COMs and DSC were calculated between each AC_fold and its respective AC_total. In the thalamus, a cluster-level inference approach (i.e. ClustFW_p001) yielded the smallest distance between COMs and highest dice scores, initially suggesting that this method may localize thalamic activation clusters more reliably. However, ClustFW_p001 also produced the highest number of folds *without* activation clusters, especially in MGN and IC regions, implying that this approach might be suboptimal when focusing on small, subcortical nuclei despite its promising stability results. Apart from cluster-based methods, the Top5%_tstat approach stands out as a strong alternative, yielding the next-best results with a mean COM distance of 2.73 mm (less than 2 voxels) and a mean DSC of 0.66, while maintaining a low number of folds without activation clusters. Thus, this approach might result in a more appropriate compromise between activation identification and stability in thalamic regions.

Similarly, within the cerebellum, the top t-statistic approaches yielded the highest accuracy for both stability metrics. Notably, while these approaches resulted in four folds lacking activation clusters, these folds only corresponded to sub-02, who exhibited predominant left cerebellar activation. As previously mentioned, left cerebellar laterality during auditory stimuli has been reported in previous work (McLachlan & Wilson, 2017; Petacchi et al., 2005), suggesting that these t-statistic approaches may still be suitable for cerebellar regions. In the midbrain, VoxFDR_p05 yielded the smallest mean COM distance and highest mean DSC between hemispheres. This method’s superior robustness in detecting activation clusters, even with lower scan duration, paired with its stability across datasets, suggests it is a highly effective approach for brainstem nuclei analysis.

Given our functional resolution (1.731 × 1.731 × 4 mm^3^), the mean difference in COM across methods was approximately 2 voxels for the thalamus and 1 voxel for the cerebellum and midbrain. Overall, these results suggest that all methods present stable localization across regions, but greater precision may be found in the identification of activation clusters in cerebellar and midbrain regions. This finding is unsurprising in the cerebellum, where a larger sized cluster (Lyu et al., 2024) is more likely to be consistently identified. Although the thalamic and brainstem auditory nuclei are similar in size (∼4x4x4 mm^3^) (Glendenning & Masterton, 1998; Sitek et al., 2019), a potential reason behind the slight discrepancy between their difference in COM may be the specific alignment of our FOV in our whole-brain acquisition. Axial slices were aligned perpendicular to the base of the fourth ventricle to better align with brainstem anatomy and maximize whole-brain coverage in our multi-echo fMRI acquisition. Although the benefits from this orientation are primarily evident in the brainstem, it is possible that this alignment may inadvertently introduce additional challenges for thalamic regions. The effects of slice orientation have been shown to impact the sensitivity of functional activation at 3T (Gustard et al., 2001), and for small thalamic nuclei, like the MGN, with diverse neuronal sizes and varying orientation in the fiber architecture of its subdivisions (Winer, 1984), these effects may be more pronounced.

Across methods, the mean DSC was 0.66, 0.82, and 0.71 for thalamus, cerebellum, and brainstem, respectively. Using similar descriptions from previous work (Wilson et al., 2017), the spatial overlap in the thalamus and midbrain can be classified as moderate-high, whereas the agreement in cerebellar regions can be classified as high, suggesting adequate reliability across methods. Recent research has discussed the limitations of the dice score in the context of reproducibility (Seghier, 2024; Soltysik, 2020). The dice score may overestimate performance for larger sized clusters whereas smaller sized clusters may yield lower scores (Seghier, 2024). Therefore, it is likely that the differences across regions may be further impacted by the size of the activation clusters. This limitation is further evidenced in our results where subjects with smaller activation clusters like sub-01 and sub-02, represented in blue and red respectively in ***Figure 8C***, consistently presented lower dice scores compared to subjects with larger activation clusters.

Furthermore, the validity of the use of this activation overlap metric has been questioned due to reports of artificial increases in dice scores from methods that may lower specificity (Soltysik, 2020). Since cluster-based approaches have been shown to present lower spatial specificity with large clusters (Woo et al., 2014), such as in the cerebellum, it is important to consider the possibility of an artificial inflation in spatial agreement in this region. Given these limitations, interpretation of results for this metric may require special consideration.

### 4.6 Recommendations for inference testing in densely sampled single-subject data

Across regions, our findings demonstrate that a whole-brain voxel-level approach with FDR correction p<0.05 (i.e. VoxFDR_p05) presented highest sensitivity in the identification of expected auditory activation, even with less data and in challenging regions like the thalamus and brainstem. Therefore, this established approach may be an effective option to analyze whole-brain task-based densely sampled individual-subject data. In the instances where a tighter delineation of the activated regions is desired, a whole-brain voxel-level approach with more conservative FDR thresholding, such as p<0.01 and p<0.001, may be considered, especially in regions such as the cerebellum. However, for systems-level studies, these more stringent thresholds may fail to identify activation in subcortical regions for individuals with smaller activation clusters. When investigating well-known systems with *a priori* knowledge of expected activated regions, a region-specific top t-statistics approach may offer the advantage of a more localized detection of the most functionally active voxels within these regions, providing more specific information regarding their predominant functional response. Additionally, this approach may be preferred in thalamic and cerebellar regions as it offers increased stability in the spatial localization and spatial overlap of the activated clusters. Given that both t-statistic thresholds (i.e. 5% and 10%) offered comparable benefits across regions, researchers may opt to use the more “liberal” option for an initial functional exploration of the desired region.

### 4.7 Limitations and Considerations

The goal of this study was to examine and compare commonly used inference testing approaches across the whole brain using a controlled, task-based precision mapping dataset. Thus, we did not comprehensively test all potential inference testing schemes. Nonetheless, the consensus in the identified activation clusters across various whole-brain and region-specific analyses provides additional confidence in the robustness of these results. Future work can focus on applying different methods and providing a wider range of statistical thresholds across inference schemes. An exciting option could be the use of Bayesian approaches (Flandin & Penny, 2007; Magerkurth et al., 2015; Penny et al., 2005; Wang et al., 2024), which have shown promising results in group and individual-level cortical studies (Mejia et al., 2022; Spencer et al., 2022).

A further limitation to consider is the earbud malfunction experienced by sub-04* during data collection. The participant only reported the malfunction at the end of the session, which precluded any technical interventions during the acquisition. Despite disruption to the auditory stimulus, expected monaural auditory activation was identified across regions. Our findings of unilateral activation in left MGN and left IC suggest a left earbud malfunction, as previous work has shown a predominant contralateral drive in auditory regions downstream of the decussation at the level of the superior olive (Devlin et al., 2003; Gutschalk & Steinmann, 2014; Mei et al., 2013; Schönwiesner et al., 2007). Furthermore, the bilateral activation present in the auditory cortex aligns with previous work involving monaural stimuli (Bauernfeind et al., 2018) and the bilateral cerebellar activation is possibly a response from cortical inputs (McLachlan & Wilson, 2017).

An additional consideration involves the inclusion of the region-specific top % t-statistic approach in this study. It is important to note that this approach does not employ any significance correction methods, and thus, a direct comparison in regard to the characterization of significance with the other inference methods cannot be achieved. Nevertheless, this technique captured the expected activation within known auditory processing regions, suggesting this approach might be a useful functional localizing tool.

Lastly, in this study, our densely sampled functional data totaled 40 minutes for each individual. While it has been previously shown that 30 minutes of data may be sufficient to identify individual-specific bilateral activation in the auditory system (Medina et al., 2026), longer scan durations may improve reliability (Lynch et al., 2020) and, consequently, affect these conclusions. In particular, our results suggest that the non-parametric permutation methods may have been limited by the amount of functional data collected in this study. Thus, a longer dataset may be necessary to evaluate and compare this approach with other methods, as well as determine if it offers any additional benefits at the individual-subject level. Nonetheless, collecting larger amounts of data may be costly, clinically impractical, and lead to decreased levels of participant task-compliance (e.g. due to drowsiness (Lynch et al., 2020; Tagliazucchi & Laufs, 2014)), meaning that approaches that are robust even with shorter scan times may be more beneficial. Furthermore, it is important to consider that the current dataset was collected with a multi-echo acquisition that prioritized in-plane resolution (1.731 × 1.731 mm^2^) with the tradeoff of a relatively large slice thickness (4 mm). Thus, these results may vary when using data with higher spatial resolution in all directions or with single-echo data, which may require longer scan times to achieve similar sensitivity (Lynch et al., 2020, 2021).

## 5. Conclusions

In this study we evaluated whole-brain and region-specific inference testing approaches with varying statistical thresholds to investigate the capacity of each to identify activation in expected regions of the auditory system and to characterize significance in densely sampled individual-subject data. The whole-brain approaches involved standard voxel-level and cluster-level inference schemes with varying statistical thresholds, as well as a non-parametric permutation inference approach. The region-specific approaches involved an exploratory top % t-statistics method and non-parametric permutation inference approaches. We compared each approach’s ability to identify activation in expected thalamic, cerebellar, and brainstem auditory regions and found that a whole-brain voxel-level approach with a false discovery rate (FDR) correction (p<0.05) presented highest sensitivity across regions and subjects in this dataset. Additionally, we tested the reproducibility and stability in the identification of these auditory activation clusters with folds of 30 minutes of data. We found that a region-specific top % t-statistic approach presented a suitable balance between activation detection and stability with low difference in each fold’s center of mass and high spatial agreement with its 40-minute data cluster (i.e. its internal benchmark) in thalamic and cerebellar regions. Although this approach does not inherently test for traditional statistical significance, it may be a useful exploratory functional localization tool and a complementary method to standard inference testing approaches. In addition, the whole-brain voxel-level approach (FDR, p<0.05) showed similar stability benefits in the brainstem and the least number of folds without activation clusters across all regions, suggesting that this approach might provide robust identification of significant bilateral activation across different regions even with lower scan duration.

## Data and code availability

Data will be made publicly available on OpenNeuro prior to publication. Code is publicly available at https://github.com/BrightLab-ANVIL/PreProc_BRAIN, https://github.com/BrightLab-ANVIL/WholeBrain_AuditoryMapping, and https://github.com/BrightLab-ANVIL/PrecisionMapping_Statistics

## CRediT authorship contribution statement

Michelle C. Medina: Writing – review & editing, Writing – original draft, Visualization, Software, Project administration, Methodology, Investigation, Formal analysis, Data curation, Conceptualization. Neha A. Reddy: Writing – review & editing, Software, Methodology, Investigation, Formal analysis, Conceptualization. Molly G. Bright: Writing – review & editing, Supervision, Project administration, Methodology, Funding acquisition, Conceptualization. Kevin R. Sitek: Writing review & editing, Supervision, Methodology, Funding acquisition, Conceptualization.

## Acknowledgements

This work was supported by the American Heart Association 25PRE1356822, NIH grants NIDCD K01DC019421, T32EB025766 and R03HD113915, the Center for Translational Imaging at Northwestern University, and through the computational resources and staff contributions provided for the Quest high performance computing facility at Northwestern University, which is jointly supported by the Office of the Provost, the Office for Research, and Northwestern University Information Technology. The authors would also like to thank Rachael Young and Megan Dorn for their support with data collection.

