## Supplemental Figure 1 for "Evaluating Approaches for Inference Testing of Whole-Brain Densely Sampled Single-Subject Task fMRI Data"

**Supplemental Material**

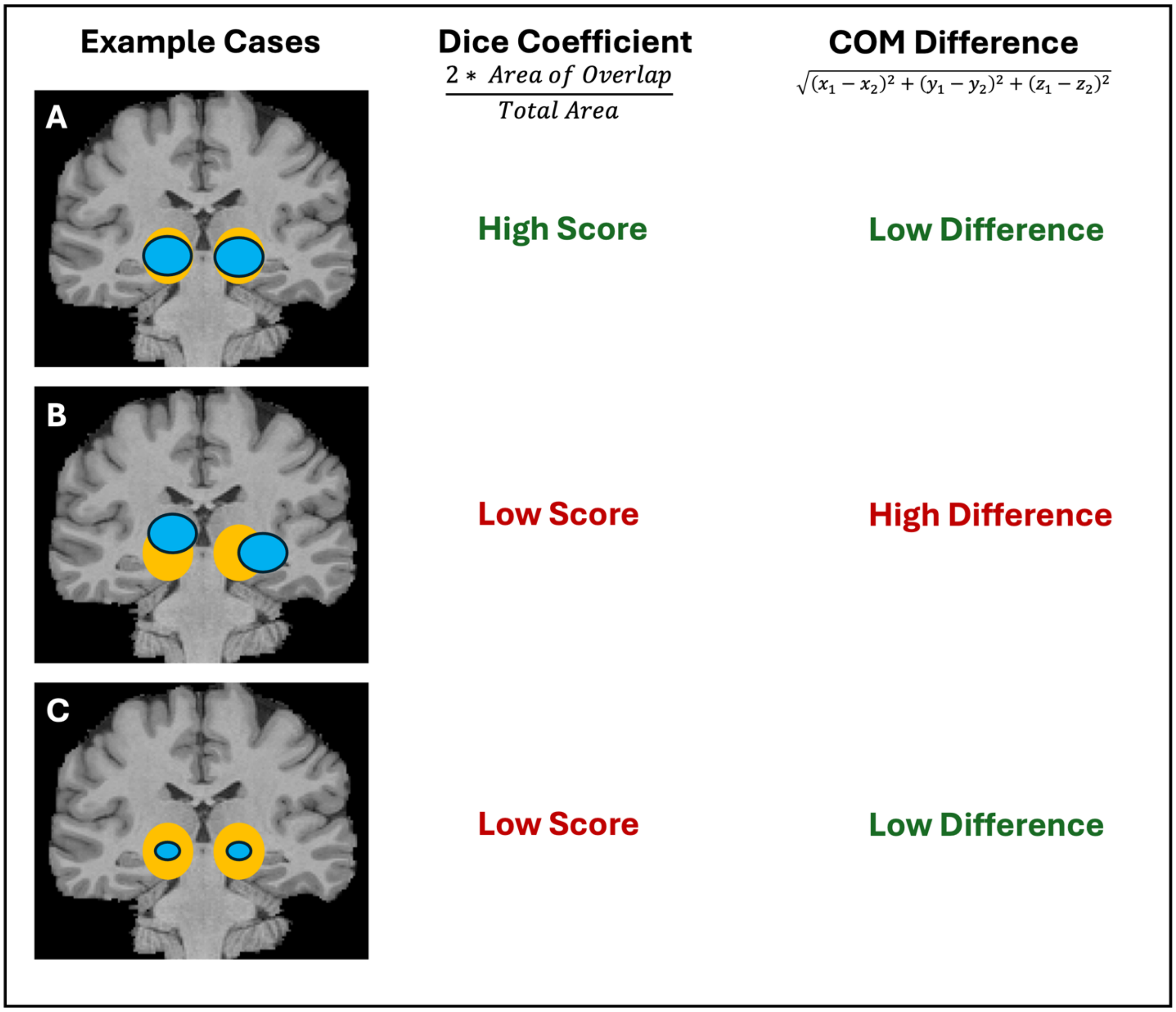


**Supplemental Figure 1:** Schematic demonstrating example cases that may yield low/high dice coefficient scores and low/high difference between center of masses, where a high dice score and low difference is preferred. The first set of activation clusters is depicted as yellow circles and second set is depicted as blue circles.
